# Telomeric Repeat-Containing RNA Increases in Aged Human Cells

**DOI:** 10.1101/2024.11.14.623513

**Authors:** Yu-Hung Hsieh, Chin-Hua Tai, Meng-Ting Yeh, Yu-Chen Chen, Po-Cheng Yang, Chien-Ping Yen, Hong-Jhih Shen, Chan-Hsien Yeh, Hung-Chih Kuo, Der-Sheng Han, Hsueh-Ping Catherine Chu

**Author notes:** To whom correspondence should be addressed. (Hsueh-Ping Catherine Chu), (Der-Sheng Han). These authors contributed equally (Yu-Hung Hsieh, Chin-Hua Tai, Meng-Ting Yeh).

## Abstract

Telomeric repeat-containing RNA (TERRA), transcribed from subtelomeric regions towards telomeric ends, poses challenges in deciphering its complete sequences. Utilizing TERRA-capture RNA-seq and Oxford Nanopore direct RNA sequencing to acquire full-length TERRA, we annotate TERRA transcription regions in the human T2T-CHM13 reference genome. TERRA transcripts encompass hundreds to over a thousand nucleotides of telomeric repeats, predominantly originating from 61-29-37 bp repeat promoters enriched with H3K4me3, RNA pol II, CTCF, and R-loops. We develop a bioinformatics tool, TERRA-QUANT, for quantifying TERRA using RNA-seq datasets and find that TERRA increases with age in blood, brain, and fibroblasts. TERRA upregulation in aged leukocytes is confirmed by RT-qPCR. Single-cell RNA-seq analysis demonstrates TERRA expression across various cell types, with upregulation observed in neurons during human embryonic stem cell differentiation. Additionally, TERRA levels are elevated in brain cells in the early stage of Alzheimer’s disease. Our study provides evidence linking TERRA to human aging and diseases.

## Introduction

Chromosome ends synthesize a heterogeneous population of long noncoding RNAs called “TERRA” (1), which are composed of the telomeric repeat sequence UUAGGG and sequences unique to the subtelomeric region of each chromosome. TERRA is transcribed by RNA polymerase II (2), and its expression is regulated by CpG-islands in the subtelomeric regions and regulated during cell cycle progression (3–5). In human cells, only a fraction of TERRA undergoes polyadenylation (6), while the majority remains non-polyadenylated and is stabilized by RALY (7), a member of the hnRNP family that interacts with both protein-coding and long non-coding RNAs (8). The regulatory regions influencing TERRA expression have been identified by the presence of three repetitive motifs, referred to 61-29-37 base pair (bp) repeat sequences, featuring a high GC content, and common to several human chromosome ends (4). The defect in DNA methyltransferase enzyme DNMT3B results in DNA hypomethylation in the subtelomeric regions and upregulation of TERRA expression (3,9–11). Studies have shown that TERRA expression can be regulated by several transcription regulators including CTCF, ATRX, NRF1, HSF1, p53, Snail1, ZNF148, ZFX, EGR1, and PLAG1 (3,12–17). TERRA can form RNA:DNA hybrids with telomeric DNA, facilitating telomere extension in telomerase-negative cancer cells using a mechanism termed “Alternative Lengthening of Telomeres” (ALT) (18–21).

TERRA exhibits binding capabilities with telomeric DNA binding proteins (22–24) and non-telomeric DNA binding proteins such as chromatin modifiers (24). It binds to both telomeric and non-telomeric chromatin, regulating telomere integrity and gene expression *in cis* and *in trans* (23–25). For instance, TERRA interacts with ATRX, a protein involved in histone H3.3 recruitment and H3K9me3 formation (26), and prevents ATRX from binding to chromatin (24,27). TERRA depletion leads to increased ATRX recruitment to repetitive sequences such as rDNA, retrotransposons, subtelomeric sequences, and telomeric repeats, as well as the regions close to transcription start sites (24,27). Furthermore, TERRA plays a crucial role in modulating DNA G-quadruplex structures around the transcription start sites, suggesting a function of TERRA beyond telomere biology (27).

Elevated TERRA levels are accompanied by telomere attrition (5,28–30) and damage (9,31,32). For instance, uncapping telomeres by TRF2 depletion upregulates TERRA expression (32). Cells derived from patients with ICF (Immunodeficiency, Centromeric instability, and Facial anomalies) syndrome display accelerated telomere shortening, premature replicative senescence, and significantly elevated levels of TERRA (9,10). Bidirectional telomeric transcription is induced by DNA damage at telomeres in Hutchinson-Gilford progeria syndrome (HGPS) fibroblasts (33). Reducing telomere length by ablating Bqt4 in fission yeast results in a greatly increased TERRA expression (34). When both telomerase and homologous recombination mechanisms are absent in budding yeast, the accumulation of RNA:DNA hybrids at telomeres results in telomere loss and increased rates of cellular senescence (35). TERRA levels are elevated at senescence in budding yeast cells lacking telomerase, while TERRA ablation by expression of an artificial antisense TERRA (anti-TERRA) delays senescence (36). Furthermore, telomere shortening induced by TNF-α in mice is mediated by TERRA elevation (37). While major progress has been made in TERRA involvement in telomere lengthening and cellular senescence in yeast, TERRA expression during human aging is poorly investigated.

Examining TERRA transcripts presents several challenges, such as unassembled genomic sequences at various chromosome ends and the lack of an annotated TERRA profile in the human genome. To address these questions, we enriched TERRA transcripts using antisense oligos to capture TERRA and conducted Nanopore direct RNA sequencing (RNA-seq). Through sequence alignment to the T2T-CHM13 genome reference, we achieve the delineation of TERRA transcription regions at the end of chromosomes. Utilizing our bioinformatics tool to quantify TERRA levels from publicly available RNA-seq datasets, we demonstrate that TERRA is upregulated during human aging. Additionally, our analyses reveal differential TERRA expression in various tissues and in individuals with human diseases.

## Materials & Methods

### Cell culture

U2OS and HeLa cells were obtained from ATCC and consistently confirmed to be free of mycoplasma contamination. Cells were cultured using Gibco Dulbecco’s Modified Eagle Medium (DMEM) with 10% fetal bovine serum, L-glutamine, and 1% penicillin/streptomycin in a 37°C incubator supplied with 5% CO_2_.

### TERRA capture for Illumina RNA-seq

To enrich TERRA, total RNA was extracted from 1×10^7^ cells using the TRIzol™ Reagent (Invitrogen™, Cat:15596018) following the manufacturer’s protocol. The total RNA was treated with DNaseI (Invitrogen™, AM2238) and supplied with Ribonucleoside vanadyl complexes (VRC, New England Biolabs, E7760S) under 37°C for 15 minutes to eliminate DNA (0.4 Unit/μl DNaseI and 10 mM VRC). The enzyme was inactivated by 5 mM of EDTA. The DNaseI-treated RNA sample was then subjected to a second round of purification using TRIzol™ reagent. For one TERRA-capture experiment, a total of 20 μg of the DNaseI-treated RNA was used. The RNA sample was dissolved in 90 μl of 6x SSC buffer (0.9M of NaCl, 0.9M of Sodium citrate). The RNA sample and 10 μl of 1 μM biotinylated TERRA antisense probe (5’-(CCCATT)_5_-3’BioTEG) were individually denatured at 70°C for 2 minutes, and then were mixed together (final: 20 μg RNA, 2x SSC buffer, 100 nM probe) for an additional 8 minutes at 70°C. The RNA-probe annealing was carried out by gradually decreasing the temperature from 70°C to 44°C, followed by incubation at 44°C for an additional 30 minutes. To capture TERRA RNA, 100 μl of streptavidin beads (MyOneC1, Invitrogen™, Cat#65002) were prepared and mixed with the sample. The mixture was then incubated on a rotating wheel at 37°C incubator for 15 minutes. The supernatant was separated from beads by a magnet and carefully removed. Beads were washed 4 times with 2x SSC buffer containing 0.1% NP40 (0.3M of NaCl, 0.3M of Sodium citrate, 0.1% NP-40) at 37°C, followed by two washes with 1x SSC buffer containing 0.1% NP40 (0.15M of NaCl, 0.15M of Sodium citrate, 0.1% NP-40), once at 37°C and once at room temperature. Finally, beads were washed once with 1x SSC buffer without NP40 (0.15M of NaCl, 0.15M of Sodium citrate) at 37°C. Washed beads were resuspended in 100 μl of nuclease-free water and incubated at 70°C to elute RNA. The elution step was repeated twice. The eluted RNA sample was concentrated by precipitating in 3× volume of 100% ethanol (300 μl), containing 0.1× volume of 3M NaOAc (40 μl, Thermo Fisher Scientific, AM9740) and 2 μl of GlycoBlue™ (Invitrogen™, AM9515) overnight at −30℃. The RNA was precipitated by centrifuge at 4℃ and 21,100 x g. The RNA pellet was washed twice with 70% EtOH and eluted in 30 μl of nuclease-free water. RNA concentration was measured using Qubit RNA HS kit (Invitrogen™, Q32852). The captured RNA was converted into cDNA, followed by paired-end Illumina sequencing and RT-qPCR.

### Illumina RNA seq library preparation

The cDNA library was constructed using the NEBNext® Ultra II Directional RNA Library Prep Kit (New England Biolabs, E7760S) according to the manufactory instruction with some modifications. The first-strand synthesis step was carried out by SuperScript IV (Thermo Fisher Scientific, Cat: 18090010). The TERRA-captured RNA sample (30 μl) was divided into two equal halves for the reverse transcription reaction. Half of the TERRA-captured sample was reverse transcribed using 2 μM random hexamers (Thermo Fisher Scientific, 48190011) and the other half using 1 μM of telomeric C-rich primer 5’-(CCCTAA)_5_-3’. These two groups are referred to as the random and specific groups, respectively. Followed by reverse transcription, second strand synthesis, end repair, adaptor ligation, and size selection. Libraries were amplified using NEBNext Ultra II Q5 Master Mix (New England Biolabs, #M0544L) with optimal cycles determined by qPCR, and size-selected twice using AMPure XP beads. Finally, the DNA was eluted in 15 μL of 0.1× TE buffer and stored at −20℃. The library size distribution was assessed using the Agilent 2100 Bioanalyzer. Each library was quantified using the NEBNext Library Quant Kit (New England Biolabs, #E7630S). The pooled libraries (fragment size 300∼500 bp) were subjected to paired-end 150 bp sequencing on Illumina MiSeq or NovaSeq platforms.

### Reverse transcription and Quantitative PCR

TERRA enrichment was assayed by RT-qPCR using cDNA. The reverse transcription was performed using random hexamers (Thermo Fisher Scientific, #48190011) and SuperScript™ IV Reverse Transcriptase (Thermo Fisher Scientific, Cat#18090200)(final: 1μM random hexamer, 400 μM dNTP, 1×SuperScript IV buffer, 8 Unit/μl SuperScript IV reverse transcriptase, 4 mM DTT, 1.6 Unit/μl RNaseOUT). The qPCR analysis was conducted using a 1:5 dilution of TERRA-enriched cDNA and a serial dilution of control cDNA, ranging from 1× to 1000× dilution (1% to 0.001% of input), to generate a standard curve. Subtelomeric primer sequences (hg38-2q, CHM13-1q, CHM13-3q, CHM13-8p, and CHM13-15q) were listed in **Supplementary Table 6**. The qPCR reaction was performed with IQ SYBR Green Supermix (Bio-Rad, Cat: 170-8882) using the CFX Real-Time PCR Detection System. The thermal cycling conditions were as follows: Initial heat activation of polymerase at 95℃ for 3 minutes, followed by 35 cycles of denaturation at 95℃ for 15 seconds, annealing and extension at 58℃ for 1 minute. Data analysis was carried out using CFX Maestro software v1.1 (Bio-Rad). For Illumina samples, the standard curve was generated from a series of dilutions from the input. TERRA RNA abundance was normalized to GAPDH. TERRA-captured samples were compared to the control sample (no capture) to calculate TERRA enrichment. For Illumina samples, TERRA enrichment ratio was calculated using the ddCt method, where the Ct value of TERRA was normalized to that of GAPDH.

### TERRA capture for Nanopore Direct RNA-seq

We harvested cells from 200 of 150 mm culture dishes for RNA extraction. The TERRA-capture procedure followed the method described previously with a few modifications. First, the RNA samples were not treated with DNase I. Second, the input RNA amount for each capture was increased to 500 μg. Third, to achieve the required RNA quantity for Nanopore direct RNA-seq, 10 captures were pooled. RNA was concentrated by precipitation with 3× volume of 100% ethanol, 1/10 × volume of 3M NaOAc (Thermo Fisher Scientific, AM9740), and 2 μl of GlycoBlue™ (Invitrogen™, AM9515) for 3 days at −20℃. 100 captures (a total of 50 mg input RNA) were required to acquire a sufficient amount of TERRA-enriched RNA for sequencing. Subsequently, RNA was precipitated by centrifugation at 4°C at 21,100 x g for 20 minutes. The pellet was washed with 70% ethanol and eluted in 20 μl of nuclease-free water. Finally, a total of 3.4 μg RNA was obtained after capture. Next, 1.5 μg of TERRA-enriched RNA was polyadenylated using E. coli Poly (A) polymerase (New England Biolabs, #M0276S) at a final concentration of 0.25 U/μl in 1X Poly(A) Polymerase reaction buffer with 1 mM dATP at 37°C for 30 minutes. To prevent RNA degradation, the reaction included 2 Unit/μl RNaseOUT, (Thermo Fisher Scientific, Cat: 10777019). The polyadenylation reaction was terminated by adding 10 mM EDTA. Polyadenylated RNA was purified using 2× sample volume of AMPureXP RNA beads (Beckman Coulter, A66514), following the manufacturer’s protocol. RNA was eluted in 20 μl of nuclease-free water. RNA concentration was measured using Qubit RNA High-Sensitivity assay (Thermo Fisher Scientific, Q32852). The fragment size was assessed using Bioanalyzer (Agilent RNA 6000 Pico Kit). TERRA-enriched RNA was subjected to Nanopore direct RNA sequencing (Cat#SQK-RNA002) with R9.4.1 flow cell (FLO-MIN106).

### Analysis of TERRA-capture Illumina RNA-seq

The fastq files were subjected to a quality filter (Q > 20), and Illumina adaptor sequences were trimmed using Trimgalore v0.6.7 (trimgalore--illumina--pair Read1.fastq Read2.fastq -o /path/to/output). The reads were aligned to the T2T-CHM13 genome using the STAR aligner v2.7.9a (star--runMode alignReads--genomeDir /path/to/T2T-CHM13--outSAMtype BAM SortedByCoordinate--outFileNamePrefix output_path/id--readFilesIn trimmed_Read1.fastq trimmed_Read2.fastq). The aligned BAM files were indexed by samtools v1.17 (samtools index - b exp.bam -o exp.bam.bai) and converted to coverage files in bigwig format by deepTools v3.3.1 bamCoverage, with RPKM as the normalization method (bamCoverage -b BAM_files -- normalizeUsing RPKM --binSize 30 -of bigwig -o output.bw). Furthermore, a comparison was performed between TERRA-captured groups and no-captured RNA-seq data (control) using log2 ratio analysis by bigwigCompare (deepTools). The coverage files (bigwig) were visualized on the Integrative Genomics Viewer (IGV) v2.16.0. TERRA transcription regions were manually assigned according to TERRA enrichment, nanopore reads, and CAGE tags at chromosome ends of the T2T-CHM13 genome. The BED and GTF files of TERRA transcription regions were used for TERRA quantification. To quantify TERRA read counts, the alignments were extracted from the BAM files using samtools v1.17 (samtools view -b -h -M -L TERRA_region.bed --fetch-pairs input.bam -o TERRA_region_alignment.bam), and filtered with MAPQ ≥ 30.

### Analysis of TERRA Nanopore direct RNA-seq

Fast5 files were processed using Guppy base caller (guppy_GPUv5.0.7) with a default quality filter of Q > 7 (guppy -i /path/to/fast5 -c rna_r9.4.1_70bps_hac.cfg -s /path/to/output_fastq). After base-calling, reads were aligned to CHM13 reference genome using the minimap2 aligner v2.22 (minimap2 -ax splice -uf -k14 /path/to/CHM13.mmi TERRA_Nanopore.fastq > TERRA_Nanopore_aligned.sam). The alignment was converted to BAM format and sorted using samtools v1.17 (samtools view -b TERRA_Nanopore_aligned.sam -o TERRA_Nanopore_aligned.bam; samtools sort TERRA_Nanopore_aligned.bam -o TERRA_Nanopore_aligned.sorted.bam). A bam index was created by samtools v1.17 (samtools index TERRA_Nanopore_aligned.sorted.bam -o TERRA_Nanopore_aligned.sorted.bam.bai). The aligned reads displayed on the genome browser were manually examined to assist in defining TERRA transcription regions. Reads aligned to TERRA transcription regions were selected using samtools v1.17 with the TERRA transcription region bed file (samtools view -b -h -M -L TERRA_region.bed TERRA_Nanopore_aligned.sorted.bam -o Nanopore_TERRA_regions_aligned.bam), and the counts of each TERRA region were quantified. For the analysis of TERRA from individual chromosome ends (Figure 1D), the unique reads were filtered based on MAPQ (≥ 1), FLAG values (FLAG = 0, or 16 for primary alignments, FLAG = 2048, or 2064 for supplementary alignments) provided by minimap2 (**Supplementary Table 5.1**). The supplementary alignments with the same read IDs were removed (**Supplementary Table 5.2**).

**Figure 1.**
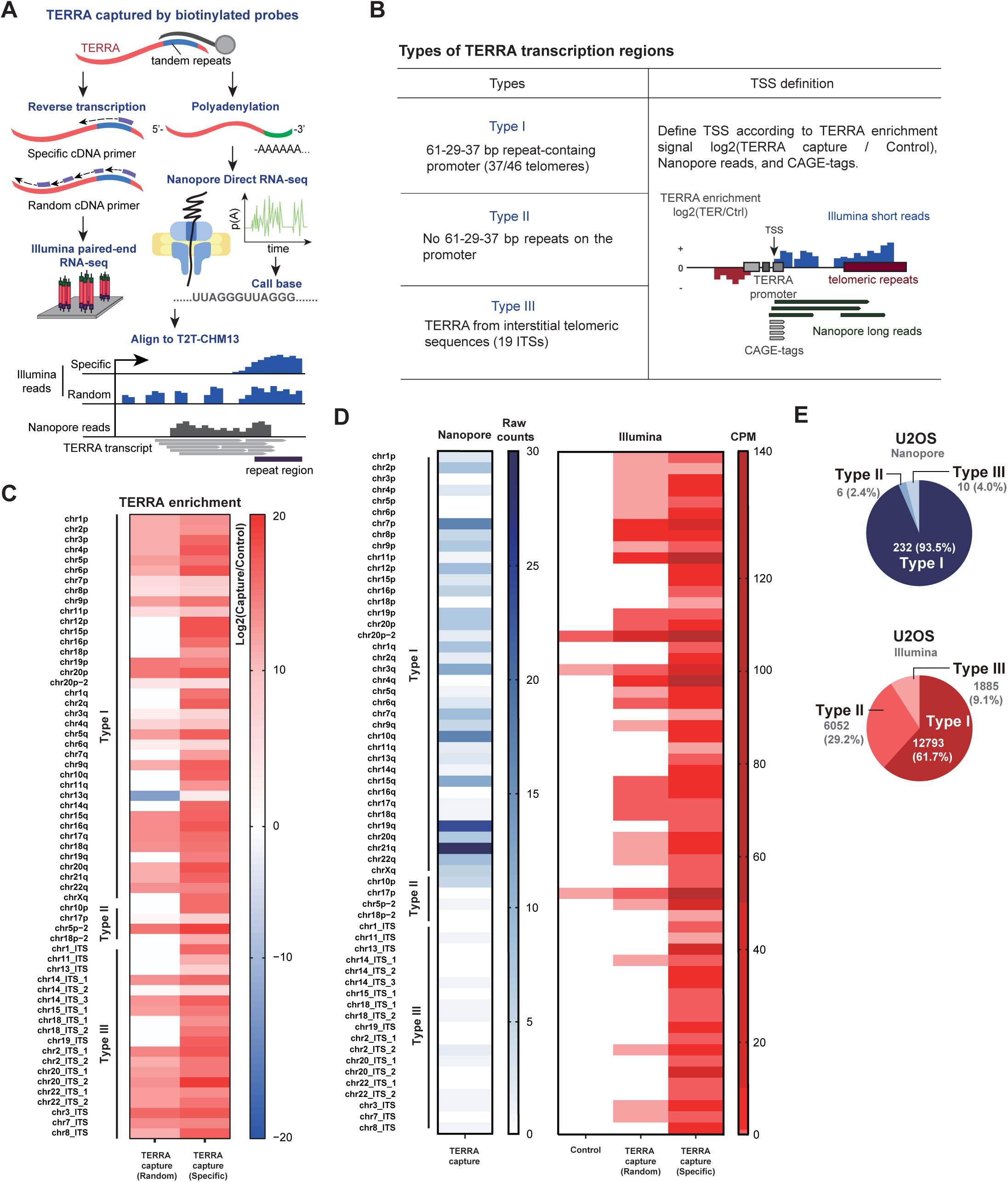
Identification of TERRA transcription regions by short-read and long-read RNA-seq in human cells. A. The flowchart of TERRA-capture RNA-seq. TERRA was captured by using biotinylated antisense oligos. Captured TERRA RNAs were subjected to Illumina or Oxford nanopore long-read RNA direct sequencing. B. Definition of three types of TERRA transcription regions. TERRA transcription start sites were determined by TERRA enrichment from Illumina short-reads, Nanopore long reads and CAGE-seq. C. Heatmap represents TERRA enrichment (log2 values) at different chromosome ends and ITSs based on Illumina read counts of TERRA transcripts in U2OS cells. Capture: TERRA capture RNA-seq. Control: regular RNA-seq with ribosomal RNA depletion. D. Heatmap represents TERRA read counts from different chromosome ends and ITSs in U2OS cells. Reads from Illumina or Nanopore sequencing were mapped to T2T-CHM13. CPM: counts per million mapped reads. E. Pie charts show the numbers of Type I, II, III TERRA reads from Nanopore or Illumina sequencing in U2OS cells.

### Analysis of ChIP-seq datasets

The ChIP-seq datasets are listed as the following: U2OS H3K4me3 (GSE114703) (38), U2OS RNA polymerase II (GSE87324) (39), U2OS CTCF (GSE87831) (40), U2OS MeDIP-seq (GSE81165) (41), U2OS DRIP-seq (GSE115957) (42), HeLa CAGE-seq (GSE121351). Additionally, the annotated CpG island BED file was downloaded from UCSC (assembly: T2T-CHM13, group: Expression and Regulation, track: CpG Islands, table: hub_3671779_cpgIslandExtUnmasked). Reads with quality scores below Q30 were filtered out, and adaptor sequences were trimmed using TrimGalore v0.6.7 (trimgalore --illumina --pair -q 30 Read1.fastq Read2.fastq -o /path/to/output). The processed reads were aligned to the CHM13 genome using bowtie2 with default settings. The resulting Sequence Alignment Map (SAM) files were converted into Binary Alignment Map (BAM) format, sorted, duplicate reads were removed, and the files were indexed using samtools v1.17 (samtools view -b exp.sam -o exp.bam, samtools sort exp.bam -o exp.sorted.bam, samtools rmdup exp.sorted.bam -o exp.sorted.rmdup.bam, samtools index exp.sorted.rmdup.bam -o exp.sorted.rmdup.bam.bai). ChIP-seq coverage files (bigwig) were generated using deepTools v3.3.1 bamCompare, normalizing using CPM method and subtracting to input signal. (bamCompare -b1 ChIP_BAM_file -b2 Input_BAM_file --normalizeUsing CPM --binSize 30 -- operation subtract -of bigwig -o output.bw). Meta-analysis surrounding the 61-29-37 nt repeat regions was performed using deepTools v3.3.1 computeMatrix and plotHeatmap (computeMatrix reference-point --referencePoint center -a 500 -b 500 -S ChIP-seq_coverage.bw -R 61_29_37_region.bed -o Matrix.gz, plotHeatmap -m Matrix.gz --colorMap bwr -o output_meta-analysis.pdf). For visualization on CHM13 genome, the bigwig files were examined in Integrative Genomics Viewer (IGV).

### Analysis of TERRA expression by TERRA-QUANT

RNA-seq datasets were downloaded and pre-processed using the previously described methods (SRAToolkit and TrimGalore commands). The processed reads were aligned to the T2T-CHM13v1.1 genome using STAR aligner with default settings. Annotated TERRA transcription regions were divided into subtelomeric and telomeric repeat portions and incorporated into the CHM13 GTF file. Read counts for both human genes and TERRA were calculated using HTseq-count (htseq -f bam -s no/yes/reverse -t transcript --idattr gene_name -m intersection-nonempty --nonunique all BAM_file CHM13_GTF_with_TERRA_regions > output_count.txt). For quantifying TERRA from individual chromosome ends, reads were extracted with MAPQ ≥ 30 for paired-end reads, or with MAPQ = 255 for single-end reads by samtools. Only reads mapped to the subtelomeres within defined TERRA transcription regions were calculated for chromosome-arm-specific TERRA. For total TERRA quantification, reads were aligned by STAR with default settings, and duplicates were removed and filtered with primary alignments (MAPQ ≥ 1) by samtools. The reads mapped to TERRA regions and all human genes were then analyzed together and normalized using the YARN package (44) in the R language. The total TERRA levels in human tissues were calculated based on reads mapped to all TERRA transcription regions with proximal TSSs located at chromosome ends (Type I and Type II, but not Type III), including both subtelomeric regions and pure telomeric repeats. “TERRA levels” refer to normalized TERRA read counts, representing the relative expression of TERRA transcripts compared to a control gene set across all samples. Counts for other genes were obtained using HTseq-count, and all counts were merged for downstream normalization. For normalization, gene counts—including TERRA and other genes quantified using HTseq-count—were processed using the YARN R package, which supports multi-sample normalization across diverse tissues and cell types.

### NGS sequencing for TERRA RT-qPCR products

RNA was extracted using TRIzol™ Reagent (Invitrogen™, Cat:15596018). Reverse transcription was performed using the SuperScript™ IV system (Thermo Fisher Scientific, Cat# 18090200) with random hexamers (2.5 nM). The RT-qPCR reaction was conducted by using IQ SYBR Green Supermix (Bio-Rad, Cat: 170-8882). The design of the primer sequences was based on the CHM13 genome (**Supplementary Table 6**). TERRA RT-qPCR products were purified using Gel/PCR Purification Mini Kit (FAVORGEN, Cat# FAGCK 001-1) before DNA library preparation. Eluted DNA was subjected to library construction using NEBNext Ultra II DNA Library Prep kit (NEB, Cat#E7645S). Paired-end 150bp reads were obtained from a Illumina NovaSeq system. The sequencing reads of TERRA RT-qPCR products underwent quality filtration (Q > 30) and adapters pruning using TrimGalore (v0.6.3) (https://github.com/FelixKrueger/TrimGalore). Trimmed reads were aligned to the CHM13 genome using STAR aligner, filtering with MAPQ ≥ 30 by samtools. SAM files were transformed to BAM files, and were deduplicated using samtools (v.1.18). Bigwig files were generated by DeepTools (v.3.3.1) bamCoverage (bamCoverage -- normalizeUsing CPM, --centerReads, and --extendReads 150). To assess the proportion of PCR products in the TERRA transcription regions, we calculated the read counts of PCR products on TERRA transcription regions using samtools (v.1.13) view (samtools view --fetch-pairs, -h, -b, -M (--use-index), and -L (--target-file)) with a bed file including TERRA transcription regions.

### Poly(A)+ and poly(A)-TERRA capture sequencing and analysis

TERRA RNA was first enriched by hybridization with a biotin-labeled C-rich telomeric probe. Following this, polyadenylated TERRA transcripts were selectively isolated using the NEBNext® Poly(A) mRNA Magnetic Isolation Module (E7490S, New England Biolabs), allowing for the separation of poly(A)+ and poly(A)− fractions. Both RNA fractions were then subjected to strand-specific library preparation (NEBNext® Ultra II Directional RNA Library Prep Kit, E7760S) for Illumina sequencing, using 150 bp paired-end reads. To further enhance the representation of TERRA transcripts during cDNA synthesis, C-rich telomeric primers were used, ensuring preferential reverse transcription of TERRA RNA. The fastq files obtained from TERRA-capture Illumina RNA-seq were subjected to a quality filter (Q > 30), and Illumina adaptor sequences were trimmed using Trimgalore v0.6.7. The reads that passed the quality filter were aligned to the T2T-CHM13 genome using the STAR aligner v2.7.9a. (star --runMode alignReads –genomeDir/path/to/T2T-CHM13 --outSAMtype BAM SortedByCoordinate --outFileNamePrefix output_path/id --readFilesIn trimmed_Read1.fastq trimmed_Read2.fastq). To quantify TERRA read counts, the alignments were extracted from the BAM files using samtools v1.17 (samtools view -b -h -q 30). Reads mapped to TERRA transcription regions including Type I, II, and III, were gathered as total TERRA reads. The proportion of TERRA reads in poly(A)+ and poly(A)-samples was calculated as the number of TERRA reads in each region divided by the total TERRA reads. The ratio of poly(A)+ to poly(A)-reads was determined by % of total TERRA reads in poly(A)+ over % of total TERRA reads in poly(A)-. Enrichment of poly(A)+ relative to poly(A)-was represented as the log_2_ values of this ratio.

### Research participants for collecting blood cells

Blood samples were collected as previously described (38). The study was approved by the institutional review board (IRB No. 201601091RIND) of the National Taiwan University Hospital. All participants were provided with written informed consent before participation in the study. Trial registration: ClinicalTrials.gov: NCT02779088. Registered 20 May 2016, https://clinicaltrials.gov/ct2/keydates/NCT02779088.

### Telomere length measurement in blood cells

Blood samples were collected from the participants as previously described (38). Briefly, genomic DNA was extracted from frozen human buffy coats using the QIAamp® DNA Mini Kit (QIAGEN, Hilden, Germany, #Cat 51306). Quantitative analysis was performed by quantitative polymerase chain reaction (qPCR). The relative telomere length for each participant was calculated using the telomere-to-single copy gene (T/S) ratio, defined as the number of telomeric repeats (T) divided by a standard reference DNA (S). The *36B4* gene was used as the single-copy DNA. A standard curve was generated from a series of dilutions of genomic DNA from HeLa cells, and was used for calibration for each qPCR plate. Primers for telomeric DNA and *36B4*: *36B4* forward primer sequence: 5ʹ-CAGCAAGTGGGAAGGTGTAATCC-3ʹ, and *36B4* reverse primer sequence: 5ʹ-CCCATTCTATCATCAACGGGTACAA-3ʹ; telomeric DNA forward primer sequence: 5ʹ-GGTTTTTGAGGGTGAGGGTGAGGGTGAGGGTGAGGGT-3ʹ and telomeric DNA reverse primer sequence: 5ʹ-TCCCGACTATCCCTATCCCTATCCCTATCCCTATCCCTA-3ʹ.

### Quantification of TERRA by RT-qPCR in blood samples

RNA was extracted from buffy coat samples using TRIzol (Thermo Fisher Scientific, Cat# 15596018) and purified by acid-phenol:chloroform (Thermo Fisher Scientific, Cat# AM9722). RNA was treated with ezDNase™ Enzyme (Thermo Fisher Scientific) to eliminate contaminating genomic DNA. Reverse transcription was carried out using the SuperScript™ IV system (Thermo Fisher Scientific, Cat# 18090200). Random (2.5 nM) hexamers combined with telomeric repeat primers (0.25 nM) to enrich TERRA complementary DNA (cDNA). The protocol for total TERRA qPCR was similar to that for the telomere qPCR using telomeric repeat primers. Beta-2 microglobulin (*B2M*) was used as the reference gene. B2M primers: forward primer sequence, 5’-CTATCCAGCGTACTCCAAAG-3’; reverse primer sequence, 5’-GAAAGACCAGTCCTTGCTGA-3’.

TERRA RNA extracted from U2OS cells was used to obtain a standard curve for calibration between samples from different qPCR plates. The relative TERRA expression level was analyzed using the TERRA RNA/B2M RNA ratio. Subtelomeric primers (hg38-2q) for TERRA RT-qPCR in blood cells were listed in **Supplementary Table 6**. Data points from samples with poor RNA quality—assessed by RT-qPCR analysis of the B2M control were excluded from the analysis.

### Analysis of TERRA expression across different tissues

Publicly available RNA-seq datasets were gathered from NCBI GEO datasets (32,39,40). Sample sources were documented in **Supplementary Table 7**. To compare TERRA levels across different tissues or cell lines, we employed the YARN R package (41). The whole gene counts derived from all RNA-seq datasets were merged in the same expression table for the following process. The filtration of low-expressed genes was performed based on an expression threshold (CPM < 1). TERRA counts for each sample were determined using TERRA-QUANT as described previously. Normalization of all data was performed using the qsmooth method (41).

### Single-cell and Single-nucleus RNA-seq analyses

Neuronal single-cell RNA-seq datasets (GSE186698) and PBMC single-cell RNA-seq datasets (GSE157007) were downloaded as previously described (**Supplementary Table 8**). Alignment, barcode assignment and Unique Molecular Identifier (UMI) counting were performed using CellRanger v7.2.0 (42) with a default setting (MAPQ = 255). Downstream analysis of CellRanger outputs was performed using Seurat v4.4.0 (43). The low-quality cells were filtered out if the number of genes detected or mitochondria gene reads were three Mean Absolute Deviations (MAD) away from the average. Genes expressed in fewer than 3 cells are excluded. 2,000 highly variable genes were identified using the variance stabilizing transformation (VST) method and were used for Principle Component Analysis (PCA). The top 20 principal components were selected for cell clustering. The result was visualized with Uniform Manifold Approximation and Projection (UMAP). Data integration was performed by using FindIntegrationAnchors() and IntegrateData() functions according to the union of the top 2,000 highly variable genes of each data. Plots of TERRA, ATRX, and MAP2 were generated by scCustomize v2.0.1 (Samuel Marsh, Maëlle Salmon, & Paul Hoffman. (2023). https://doi.org/10.5281/zenodo.10161832). In neuronal differentiation samples, cells positive for CD29, CD44, and CD105 but negative for CD14, CD34, and CD45 were classified as mesenchymal stem cells; cells expressing SOX2 and Nestin were identified as neuroepithelial cells, while those positive for MAP2 were categorized as mature neurons. Cells in PBMC were annotated by the expression of representative cell markers: CD4+ T cells (CD4, IL7R and CD3G), NK cells (NCAM1 and NKG7), CD8+ T cells (CD8A and CD8B), B cells (CD19 and MS4A1), Monocytes (CD14 and CD16), Dendritic cells(IL3RA). Single-nucleus RNA-seq datasets were obtained from The Rush Alzheimer’s Disease Center (RADC) Research Resource Sharing Hub at Synapse (https://www.synapse.org/#!Synapse:syn18485175) with the DOI 10.7303/syn18485175. The ROSMAP metadata can be found at https://www.synapse.org/#!Synapse:syn3157322. Details of individual samples, including those with no pathology (13 individuals), AD-early pathology (8 individuals), and AD-late pathology (5 individuals), utilized in the analysis of this study are outlined in **Supplementary Table 9**. In Alzheimer’s disease samples, we assigned cell types by marker genes: excitatory neurons (NRGN), inhibitory neurons (GAD1), astrocytes (AQP4), oligodendrocytes (MBP), microglia (CSF1R and CD14) and oligodendrocyte progenitor cells (VCAN).

### Alzheimer’s disease patient-derived iPSC differentiation to neurons

To generate cortical neurons, induced pluripotent stem cells (iPSCs) derived from an Alzheimer’s disease (AD) patient carrying the P117L mutation in presenilin-1 (*PS1*)—the catalytic subunit of γ-secretase—and mutation-corrected isogenic controls were used. iPSCs were dissociated with dispase and cultured in suspension using human embryonic stem cell (hESC) medium lacking bFGF, but supplemented with 100 nM LDN193189 and 10 µM SB431542 to initiate neural induction. Cells were then transferred to N2 medium and cultured until neural rosette structures formed. These neural rosettes were manually isolated and further cultured in NI medium to promote neural sphere formation. The resulting neural spheres were plated onto Matrigel-coated 6-well plates and cultured in ND medium to induce cortical neuron differentiation. The differentiated cortical neurons were subsequently collected for RT-qPCR analysis.

## Results

### Nanopore Direct RNA sequencing for TERRA transcripts

To investigate TERRA transcripts in human cells, we captured RNA molecules containing UUAGGG repeats by using biotinylated antisense oligos in U2OS cells (**Figure 1A**). The TERRA-enriched RNA samples were subjected to two different sequencing methods: Illumina paired-end sequencing and Nanopore direct RNA sequencing. For Illumina sequencing, TERRA-enriched RNA was reverse transcribed using either random hexamers (random group) or telomere-specific primers (specific group) for cDNA library construction. For Nanopore Direct RNA sequencing, enriched TERRA molecules were polyadenylated by poly(A) polymerase before library preparation (**Figure 1A**). Approximately, a few hundred to over two-thousand-fold enrichment was observed after TERRA capture (**Supplementary Figure S1A**). The reads from both methods were aligned to the T2T-CHM13 human genome reference, which includes nearly complete coverage of the entire genome including subtelomeric and telomeric regions (44).

### TERRA transcription regions at chromosome ends

The criteria for identification of TERRA transcription regions included the observations of Nanopore reads, CAGE (Cap Analysis of Gene Expression) tags, and TERRA enrichment within 100 kb of chromosome ends (**Figure 1B, Supplementary Figure S1B**). Compared to ribosomal depleted RNA-seq (control), TERRA-capture RNA-seq (enriched by antisense oligos) exhibited an increase in read counts derived from chromosome ends in U2OS cells, shown by TERRA enrichment in log2 values from Illumina sequencing data (**Figure 1C**). In cases where no CAGE-seq tags mapped to the subtelomeric regions, we estimated transcription start sites (TSSs) based on subtelomeric regions with Nanopore long reads and TERRA enrichment. We identified 39 chromosome ends that contain TERRA transcription (**Supplementary Tables 1, 2**). Notably, chr5p, chr18p, and chr20p each potentially contain two TERRA transcription start sites (one proximal and one distal to telomere), while other chromosome ends have a single TERRA transcription start site located near the telomeric repeat tracts (**Supplementary Figure S1C**). The distance of the identified TERRA TSS from the telomeric repeat tract per chromosome end is shown in **Supplementary Figure S2**. These results demonstrate the presence of TERRA transcription across a majority of chromosome termini in U2OS cells (**Figure 1D**), aligning with a recent study showing that TERRA is detectable at most chromosome ends in human cell lines (45).

To further verify the TERRA transcription regions, we performed TERRA-capture RNA-seq in another cell line (HeLa). The TERRA enrichment was also observed in defined TERRA transcription regions at chromosome ends in HeLa cells (**Supplementary Figure S3**), showing the consistency of TERRA transcription regions between two different cell lines.

Notably, most CAGE tags were mapped to the 37 bp repeat regions (**Supplementary Figure S4**). In agreement with this, TERRA nanopore reads were aligned with the 37 bp repeats at their 5’ termini (**Supplementary Figure S4**). These results suggest that the regions preceding the 37 bp repeats may function as potential TERRA promoters, supporting previous studies that showed the importance of the 29 bp repeat regions in regulating transcription activity (4,21).

We classified three distinct types of TERRA transcription regions according to their locations and promoters (**Figure 1B**). The majority of telomeres (37 out of 46 chromosome ends) were characterized as Type I TERRA transcription regions by the presence of 61-29-37 bp repeats at the promoter (**Supplementary Figure S4**). Only chr10p and chr17p termini were characterized as Type II TERRA transcription region, which lacks 61-29-37 bp repeats (**Figure 1B, Supplementary Figure S4**). TERRA transcription regions with 61-29-37 bp repeats or CAGE tags were summarized in **Supplementary Tables 1 and 2**. Not all Type I TERRA promoters encompass all three repetitive motifs. 16 chromosome ends are composed of all three repetitive motifs, while 21 chromosome ends lack either one or two repetitive motifs (**Supplementary Figure S5A**, **Supplementary Table 3**). To investigate the impact of the 29 bp repeat element on TERRA expression, we compared TERRA levels at chromosome ends with or without the 29 bp repeats. Our analysis revealed no significant difference in TERRA expression between these regions (**Supplementary Figure S5B**), implying the presence of an additional promoter that lacks the 29 bp repeats for TERRA transcription.

### TERRA transcription from interstitial telomeric repeats

In addition to TERRA transcription at chromosome ends, TERRA can be transcribed from interstitial telomeric sequences (ITS), classified as Type III TERRA transcription regions (**Figures 1B-1D**, **Supplementary Figure S4, Supplementary Table 4**). We quantified Type I, II, and III TERRA transcription and found that most TERRA transcripts in U2OS cells originate from chromosome ends (Type I + II) (**Figure 1E**, **Supplementary Figure S3B**). Type III accounts for only ∼4% of Nanopore and 9.1% of Illumina TERRA reads in U2OS but is more prominent in HeLa cells, comprising 46% of Nanopore and 17.7% of Illumina reads. In the T2T-CHM13 genome, we identified 79 ITS sites containing telomeric repeats longer than 200 base pairs (**Supplementary Figure S5C**). Among these, 19 sites showed TERRA enrichment and Nanopore reads containing long telomeric repeats at the 3’ ends. Notably, Type III TERRA transcription regions exhibit CAGE tags, and the promoters usually comprise the 37 bp repeats without 29 bp repeats (**Supplementary Figure S4)**. These results indicate that most TERRA transcripts originated from the 37 bp repeat regions, regardless of whether TERRA derived from chromosome ends or ITS. Genome browser views demonstrated some examples of TERRA transcription regions carrying these features (**Supplementary Figure S4**).

Interestingly, we observed antisense TERRA transcripts, named ARIA (46) in Nanopore direct RNA-seq data. ARIA reads were mapped to chromosome 2, a region containing an ancestral telomere-telomere fusion site (54). The 5’ ends of ARIA reads comprise C-rich telomeric repeats near the junction of the telomere-telomere fusion site, following the unique sequences aligned to chromosome 2 (**Supplementary Figure S5D**). TERRA transcripts were also observed near the junction but were not directly connected to ARIA transcripts (**Supplementary Figure S5D**).

### The length of telomeric repeats in TERRA transcripts

We sought to elucidate the length of telomeric repeat tracts within TERRA molecules by analyzing Nanopore sequencing reads that aligned to subtelomeric regions and extended into the telomeric repeats. Among TERRA reads mapping to chromosome ends, the maximum lengths are 2478 bp in U2OS cells and 4639 bp in HeLa cells (**Figures 2A-2B)**. The mean lengths of bulk TERRA reads were 721 bp in U2OS cells and 982 bp in HeLa cells. For the pure telomeric tracts in TERRA reads, the maximum lengths observed were 1080 bp in U2OS cells and 1487 bp in HeLa cells, while the mean lengths were 222 bp and 284 bp in U2OS and HeLa cells, respectively. Overall, telomeric repeat tracts within TERRA reads ranged from several hundred nucleotides to over one thousand nucleotides. Dot plots of Nanopore TERRA reads per chromosome arm, grouped by types of TERRA transcription regions, revealed considerable variability in length of bulk TERRA reads and telomeric repeats across chromosomes in both U2OS and HeLa cells (**Figure 2C**). The maximum bulk TERRA read lengths were longer in Type I compared to Type II and Type III in both cell cells. Mean telomeric repeat lengths in TERRA reads were slightly shorter in Type III in both cell lines. Given the fragility of RNA molecules, it is possible that some TERRA transcripts were partially degraded during sample preparation, potentially resulting in shorter observed lengths and increased variation in transcript size. Nevertheless, our findings are consistent with previous reports indicating that TERRA transcripts generally exceed 1kb in length and that the average telomeric repeat length within TERRA is approximately 200∼300 bp (32,47).

**Figure 2.**
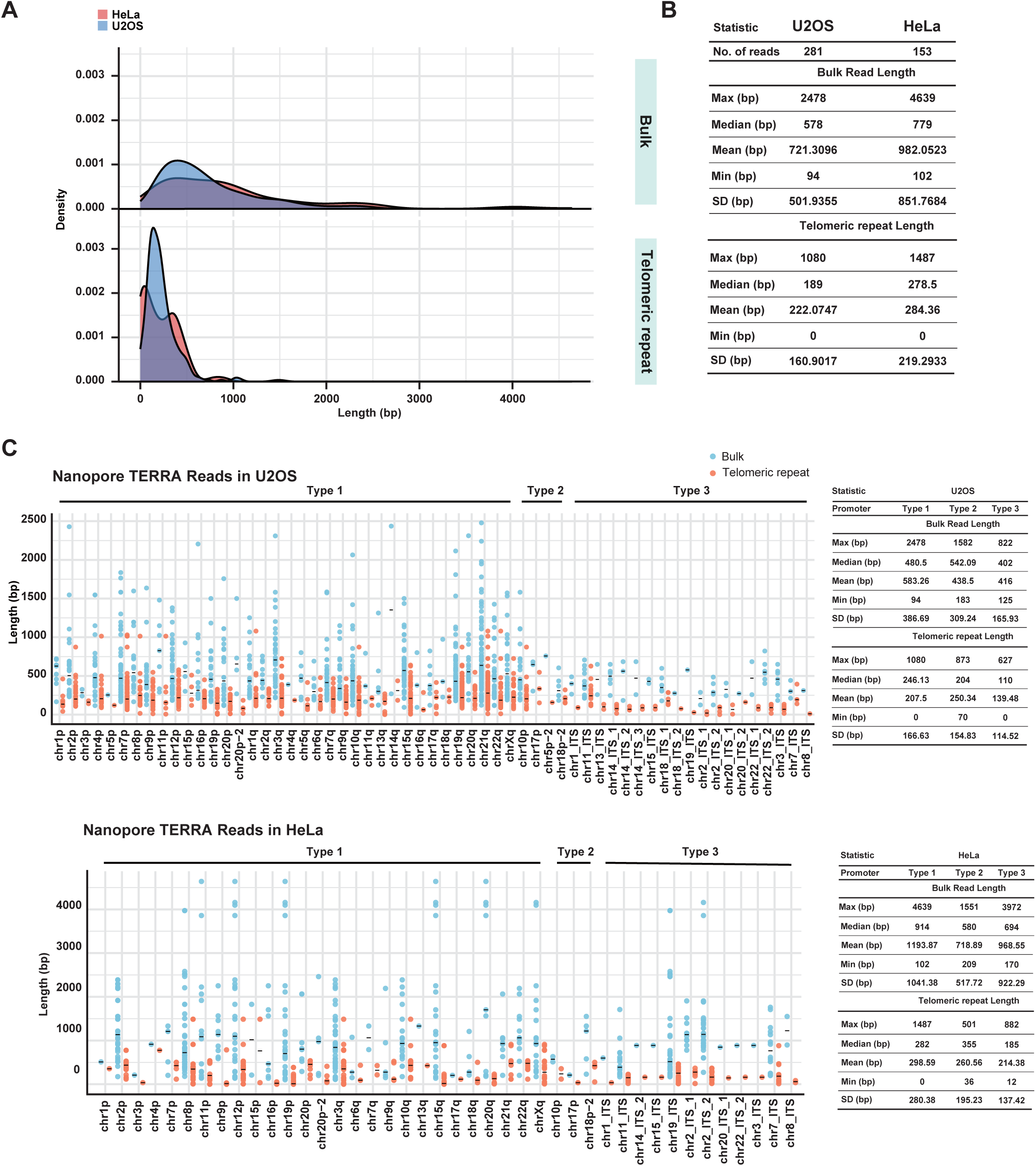
TERRA length distribution from nanopore reads in U2OS and HeLa cells. A. Density plots showing the distribution of TERRA bulk read lengths and telomeric repeat lengths in TERRA reads in U2OS and HeLa cells. Nanopore reads mapped to T2T-CHM13 TERRA transcription regions were calculated. B. Statistic analysis of TERRA bulk read lengths and telomeric repeat lengths in TERRA reads. C. Dotplots show TERRA bulk read lengths and telomeric repeat lengths in TERRA reads at each chromosome end and ITS. Each dot represents each nanopore read. Solid bar, median.

### TERRA promoters are associated with H3K4me3, Pol II, CpGs, and R-loops at distinct repeat elements in Type I TERRA transcription regions

Next, we searched for common features among the transcription start sites of TERRA. ChIP-seq data for RNA polymerase II (48), H3K4me3 (49), CTCF (50) and DNA methylation (51) were analyzed using T2T-CHM13 reference genome. Additionally, DRIP-seq data (52) was investigated to examine the profiles of R-loops surrounding the 61-29-37 bp repeats of TERRA promoters located in the subtelomeric regions of the Type I TERRA transcription regions. The meta-analysis revealed epigenetic profiles corresponding to each repeat element (**Figure 3A**). The 61 bp repeat element exhibited an enrichment of CTCF, RNA polymerase II, and R loops. The 29 bp repeat element showed an enrichment of DNA methylation compared to other repeats. Remarkably, CAGE tags and H3K4me3 were enriched at the 37 bp repeat regions. The distribution of analyzed epigenetic marks along the TERRA promoter was illustrated in **Figure 3B**, and an example of the genomic view of these marks was shown in **Figure 3C**. We conclude that DNA methylation predominantly occurs at 29 bp repeats, while RNA polymerase II and H3K4me3 are enriched at 61 bp and 37 bp repeats respectively in Type I TERRA transcription regions.

**Figure 3.**
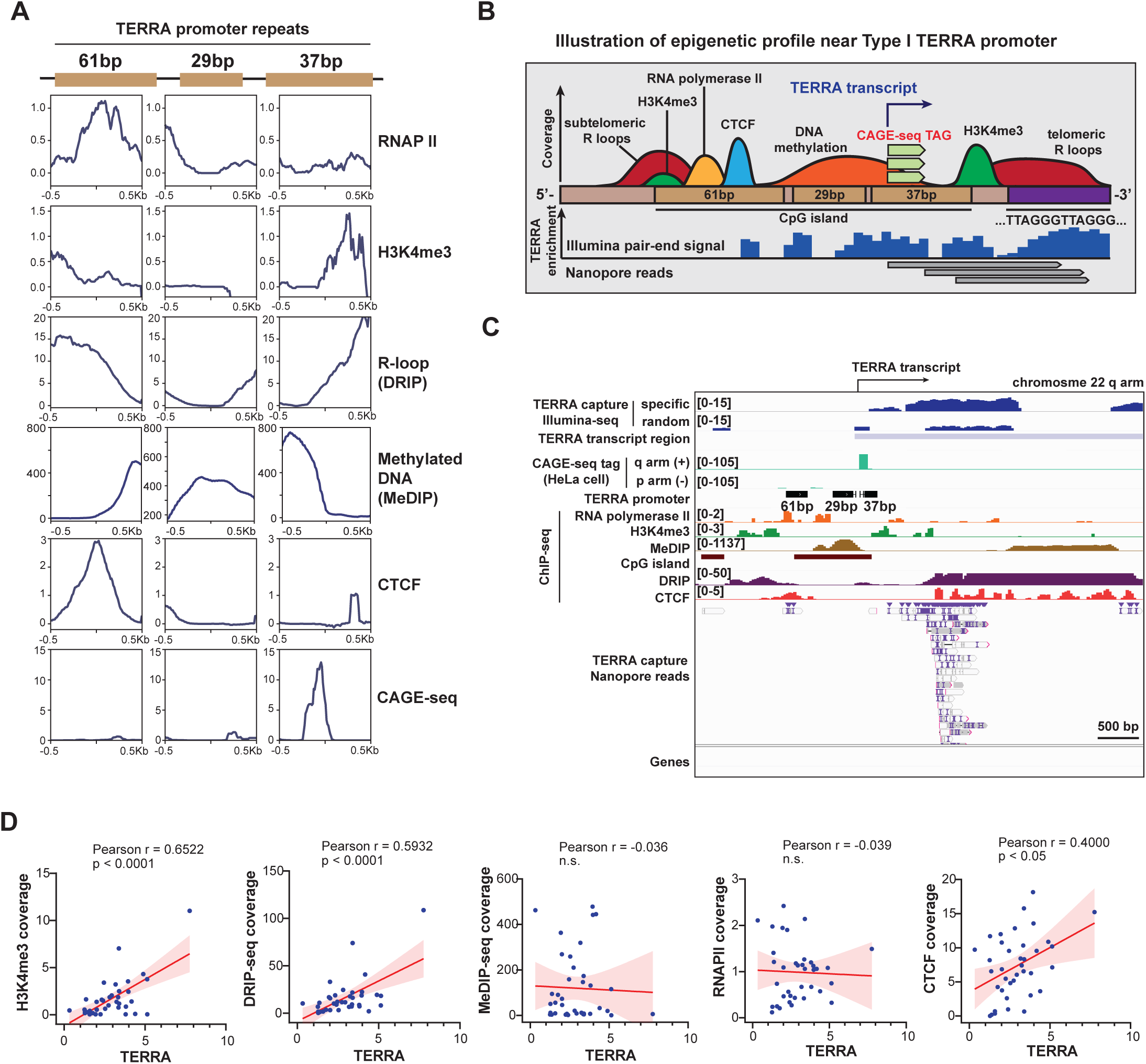
Enrichments of H3K4me3, RNA Pol II, CpGs, DNA methylation and R-loop at TERRA promoters. A. Meta-analysis showing the distribution of epigenetic marks on 61-29-37 bp repeats located at subtelomeric regions (Type I promoter). Each plot represents the coverage of indicated marks. B. A schematic model of the epigenetic profiles at Type I TERRA promoter near telomeres. C. Genome browser view showing the coverage of indicated epigenetic marks at TERRA promoter on chr22q arm. D. Scatter plots of TERRA v.s. H3K4me3, R-loop (DRIP-seq), DNA methylation (MeDIP-seq), CTCF, or RNAPII near TERRA transcription start sites. Each dot indicates TERRA enrichment and epigenetic marks at individual chromosome ends (Type I +Type II TERRA transcription regions) in U2OS cells. TERRA enrichment (log2 ratio) was calculated by comparing TERRA capture and no capture. P values, by Pearson’s correlation.

Our observations suggest that TERRA are transcribed mainly from the 61-29-37 bp repeat regions. Only chr10p and chr17p do not contain the 61-29-37 repeats adjacent to TERRA transcription regions. It’s worth noting that while certain TERRA transcription regions lack 29 bp repeats, they still consist of 61-37 bp repeats. Intriguingly, even in regions without 29 bp repeats, DNA methylation was still detected in the CpG islands between the 61 and 37 bp repeat tracts (**Supplementary Figure S6A**).

To investigate the correlation between these epigenetic marks and TERRA levels, we compared TERRA expression with the abundance of these epigenetic marks at TERRA promoters. The promoter regions were selected from 3 kb upstream of the transcription start sites, where the 61-29-37 bp repeats were located. Scatter plots demonstrated that TERRA expression is positively correlated with H3K4me3, R-loops, and CTCF at chromosome ends (**Figure 3D**).

### TERRA-QUANT for measuring TERRA expression

To analyze TERRA expression using publicly available short-read RNA-seq datasets, we established a bioinformatics tool, dubbed TERRA-QUANT, for measuring TERRA levels based on the transcription regions defined in this study (**Figure 4A**). TERRA-QUANT quantifies reads mapped to TERRA transcription regions at each chromosome end, including reads from Type I, II, and III. For total TERRA levels from chromosome ends, only TERRA reads mapped to Type I and II TERRA transcription regions were selected and subsequently underwent YARN normalization. For chromosome-end-specific TERRA analysis, we separated reads from subtelomeric regions and pure telomeric repeat tracts, and only quantified subtelomeric reads and excluded pure telomeric repeat reads.

**Figure 4.**
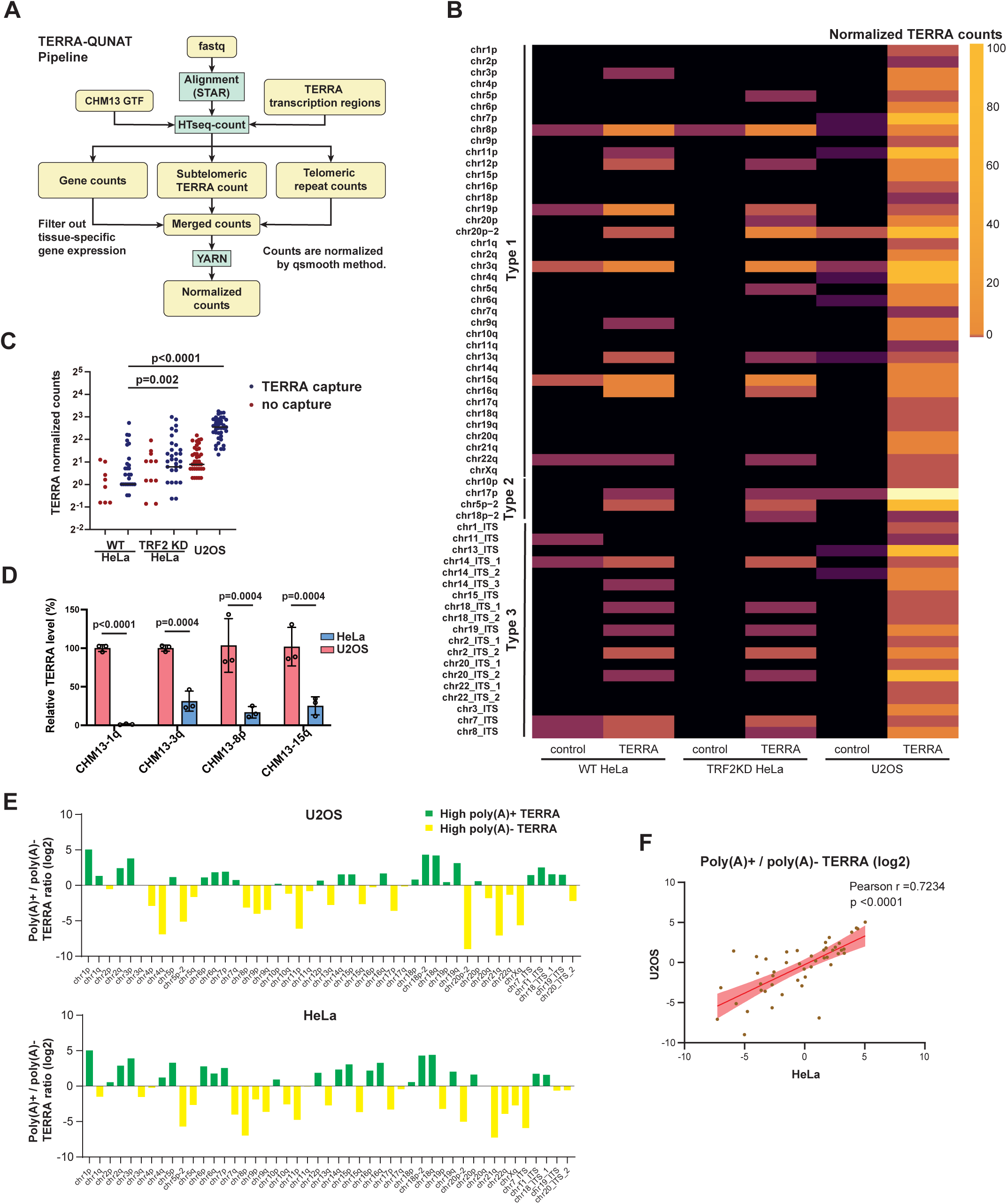
TERRA-QUANT for measuring TERRA expression. A. The workflow of bioinformatics pipeline for quantification of TERRA expression. Reads that mapped to TERRA transcription regions were counted. YARN was used for normalization between different tissues. B. Heatmap showing TERRA expression from individual chromosome ends and ITSs in HeLa and U2OS cells. TERRA capture, using antisense oligos to enrich TERRA. Control, no TERRA capture. C. TERRA normalized counts were analyzed by TERRA-QUANT. Each dot indicates TERRA counts at individual chromosome ends and ITSs. TRF2 deletion (TRF2KO) in HeLa cells increases TERRA expression. P values, by Wilcoxon matched-pairs signed rank test. Bars, median. D. RT-qPCR to detect TERRA in HeLa and U2OS cells. Subtelomeric primers for TERRA were designed based on the T2T-CHM13 genome sequence. Data from three biological replicates. P values, by two-tailed Student’s t-test. Bars, mean±SD. E. Log2 ratios of poly(A)+ to poly(A)-TERRA reads mapped to individual chromosome ends and ITSs in U2OS and HeLa cells. F. Scatter plots showing the correlation between U2OS and HeLa cells in the poly(A)+/poly(A)-TERRA ratios. Each dot represents the ratio (log2) of poly(A)+/poly(A)-reads at each TERRA transcription region.

To test the reliability of this tool, we applied TERRA-QUANT to analyze TERRA expression in HeLa cells using Illumina sequencing data obtained from a previous study (32). The previous study enriched TERRA RNA by using biotinylated antisense oligos to capture TERRA from nuclear RNA. Notably, TERRA read counts were detected at the majority of chromosome ends in the TERRA capture group, whereas they were absent at multiple chromosome ends in the control group without TERRA capture (**Figures 4B**). TRF2 deficiency led to increased TERRA read counts at multiple chromosome ends compared to wildtype cells (**Figures 4B**). Additionally, TERRA read counts from chromosome ends in HeLa cells were significantly lower than those in U2OS cells (**Figure 4C**). These results are consistent with earlier studies showing elevated TERRA levels upon TRF2 depletion (4,32), affirming the robustness of the TERRA-QUANT methodology.

To further validate the annotations for TERRA transcription regions, we designed subtelomeric primer sets specific to CHM13-1q, 3q, 8p, and 15q chromosome ends based on the T2T-CHM13 genome sequence (**Supplementary Table 6**). We used these subtelomeric primers to detect TERRA following TERRA capture and observed over 1000-fold enrichment (**Supplementary Figure S6B)**, indicating that the primers efficiently amplified TERRA transcripts. Notably, RT-qPCR analysis using these subtelomeric primers revealed increased TERRA levels from various chromosome ends in U2OS compared to HeLa cells (**Figure 4D**). Subsequently, the RT-qPCR products were subjected to Illumina sequencing to validate their sequences. We observed that the reads of RT-qPCR products were predominantly mapped to their target chromosome ends (**Supplementary Figure S6C**). These results indicate the specificity of these subtelomeric primers.

### Poly(A)+ and non-poly(A) TERRA capture sequencing

Since most publicly available RNA-seq datasets are generated from polyadenylated RNA--and only a portion of TERRA transcripts are polyadenylated--we sought to investigate the genomic origins of TERRA molecules with and without poly(A) tails. To achieve this, we performed TERRA capture followed by an additional poly(A) selection step to separate poly(A)+ and poly(A)-fractions. Both poly(A)+ and poly(A)-TERRA fractions were subjected to Illumina sequencing. We analyzed the ratio of poly(A)+ to poly(A)-TERRA reads in both U2OS and HeLa cells and found that certain chromosome ends, such as chr1p, chr2q, chr3p, and 18q, produced more poly(A)+ TERRA transcripts in both cell lines (New **Figure 4E**; **Supplementary Figure S6D**). In contrast, other chromosome ends, including chr8p, chr11p, chr15q, and chr21q, produced more poly(A)-TERRA transcripts. Notably, we observed a strong positive correlation (Pearson r=0.7, p<0.0001) between U2OS and HeLa cells in the ratio of poly(A)+ to poly(A)-across different chromosome ends (**Figure 4F**), suggesting that polyadenylation patterns in TERRA are largely consistent across cell types.

### ALT positive osteosarcomas show elevated TERRA levels

We gathered rRNA-depleted RNA-seq datasets from a previous study, which included three biological replicates of 13 human osteosarcoma cell lines, with 4 cell lines categorized as ALT negative (n=12) and 9 cell lines classified as ALT positive (n=27) (39). The RNA-seq reads of these datasets were aligned to the T2T-CHM13 genome and processed using TERRA-QUANT. Total TERRA levels were then determined by cumulating TERRA reads from all chromosome ends. Notably, ALT-positive cells exhibited a higher level of total TERRA, and a lower RNA level of telomerase reverse transcriptase (TERT) (**Figures 5A-5B**). These results agree with the previous finding showing ALT cells with elevated TERRA expression (2,47).

**Figure 5.**
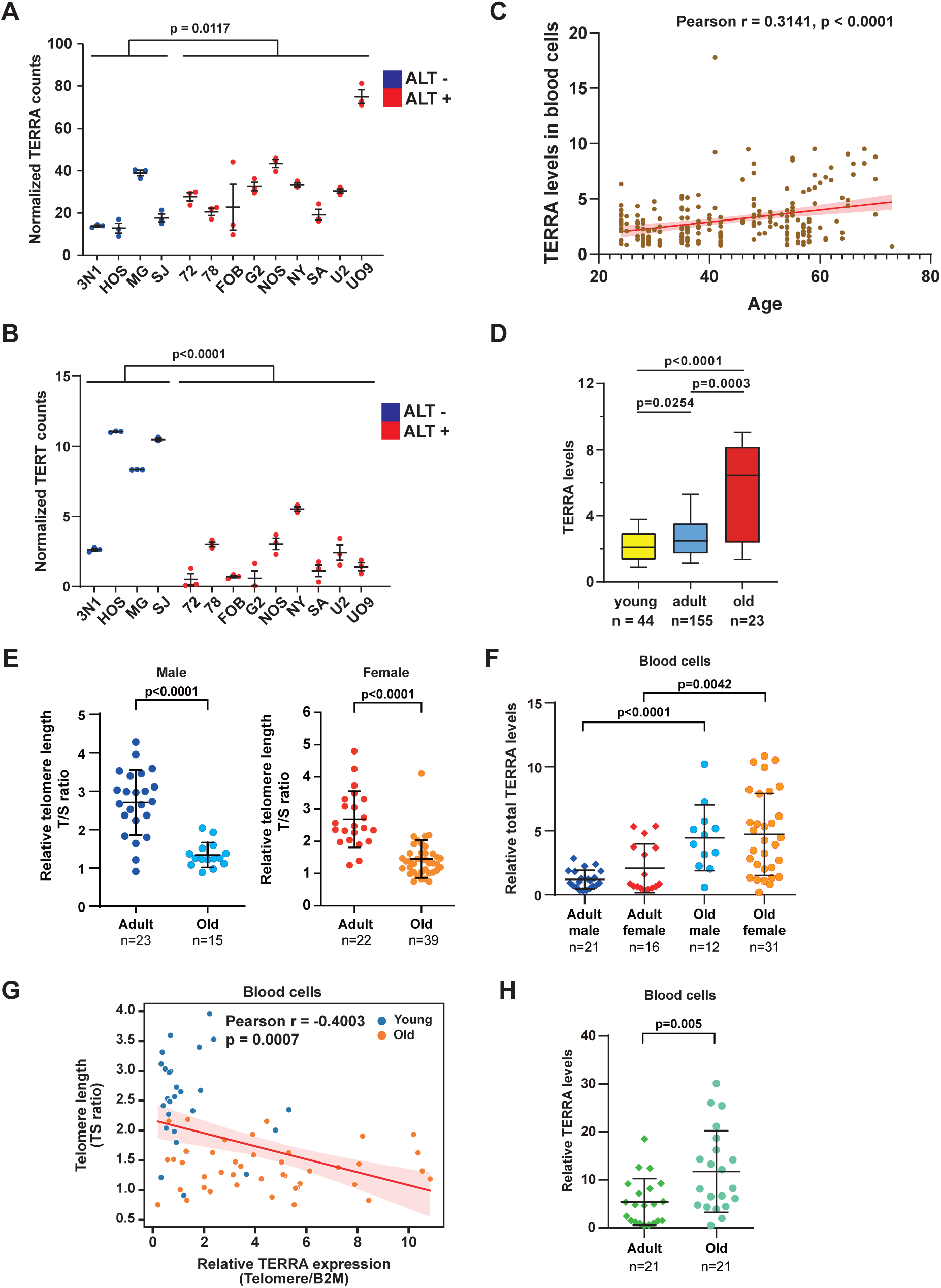
TERRA levels increase with age in blood cells. A. TERRA expression in ALT positive (+) or negative (-) cell lines was quantified by TERRA-QUANT. Total TERRA counts at all chromosome ends were accumulated. Each dot indicates the normalized TERRA read counts of each RNA-seq dataset. Bars, mean±SD. P values, by Mann-Whitney U test. B. TERT expression of each RNA-seq data from ALT+ or ALT-cells. Bars, mean±SD. P values, by Mann-Whitney U test. C. TERRA expression in human blood cells was analyzed by TERRA-QUANT using RNA-seq datasets. Scatter plots showing the correlation of TERRA v.s. age in blood. Each dot indicates the total TERRA normalized reads from all chromosome ends of each individual. P values, by Pearson’s correlation. D. Normalized TERRA counts in blood cells of different ages. Young (<30 years); adult (30∼59 years); old (≥60 years). Bars, median with interquartile. P values, by Mann-Whitney U test. E. The T/S ratio represents the relative telomere length (T) to the single-copy gene (S, 36B4 gene). Each dot represents an individual T/S ratio. Adult (21∼59 years); old (≥60 years). F. RT-qPCR to detect total TERRA levels in blood cells using telomeric repeat primers. Each dot indicates each individual. G. Scatter plot showing negative correlation between TERRA and telomere length in blood cells. P values, by Pearson’s correlation. H. RT-qPCR to detect TERRA levels using subtelomeric primers (hg38-2q). Each dot indicates each individual. (E, F and H) P values, by two-tailed Student’s t-test. n=sample size. Error bar, SD.

### TERRA increases along with human aging in blood cells

To compare TERRA expression across different ages, we collected poly(A)-enriched RNA-seq datasets from a previous study (47), which comprised 222 human blood samples for analysis. These blood samples cover a range of ages from 24 to 73 years old and were separated into three groups: young (<30 years old), mid-age (30-59 years old), and old groups (≥60 years old). Interestingly, TERRA-QUANT analysis demonstrated that total TERRA levels accumulated from all chromosome ends were positively correlated with age (**Figure 5C**). When grouped by different ages, total TERRA levels were significantly increased in old individuals, compared to young and mid-age individuals (**Figure 5D**). For chromosome-end-specific analysis, reads mapping to individual chromosome ends were low, preventing statistical analysis (**Supplementary Figure S7A**). Notably, TERRA reads were predominately mapped to Type I transcription regions (**Supplementary Figure S7B**).

Next, we confirmed the association of TERRA levels with aging by collecting blood samples from individuals of various ages and conducted quantitative PCR to measure telomere length and TERRA levels. The samples were grouped by adult (21-60y) or old (> 60y) ages. Genomic DNA was extracted from buffy coats, and telomere length (T/S ratio) was analyzed by quantitative PCR. We observed that telomere lengths significantly decreased in elderly people in both males and females (**Figure 5E**). Interestingly, total TERRA RNA levels in blood samples determined by RT-qPCR were significantly elevated in old males and females (**Figure 5F**), and were negatively correlated with telomere lengths (**Figure 5G**). Using subtelomeric primers for RT-qPCR also showed a significant increase in TERRA expression in the old group compared to the adult group (**Figure 5H**), confirming an elevation in TERRA levels in aged leukocytes.

### TERRA is upregulated in the aged brain

To explore the relationship between TERRA and human aging, we analyzed published poly(A)-enriched RNA-seq datasets obtained from various human tissues including brains, hearts, and ovaries (40) by TERRA-QUANT. The scatter plots displayed a significant positive correlation between total TERRA levels and age in brains, but not in ovaries or hearts (**Figure 6A**). Remarkably, total TERRA levels in brain tissues exhibited an upregulation in the old group (≥60 years old), compared to the adult group (30-59 years old) (**Figure 6B**). Considering that ATRX is important for neuronal development and functionality (53–55), we analyzed ATRX expression in human brains. Interestingly, the levels of ATRX, which is a TERRA-interacting protein (24), declined in old individuals (**Supplementary Figure S7C**) and displayed an anticorrelation with TERRA expression in brain tissues (**Supplementary Figure S7D**).

**Figure 6.**
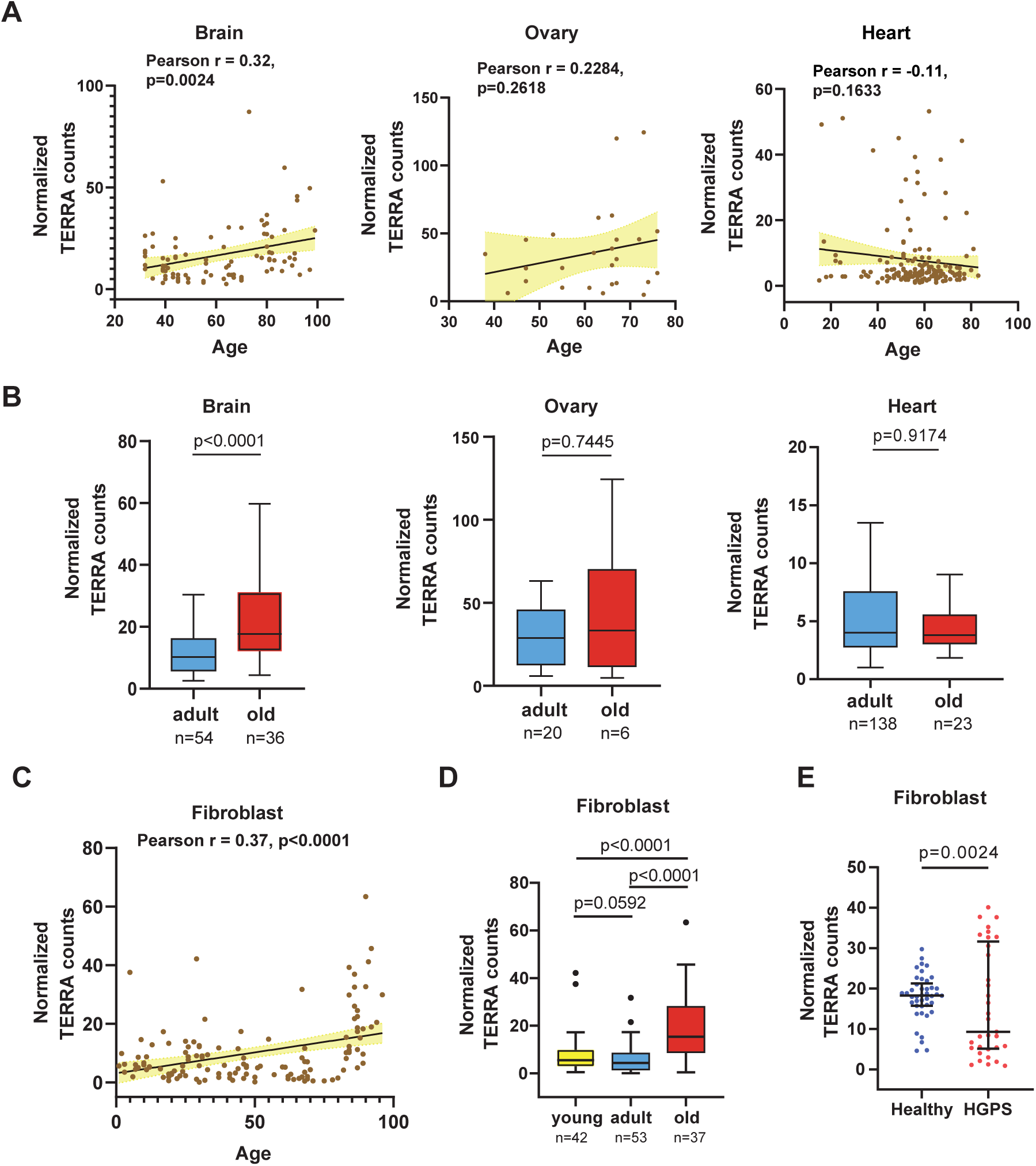
TERRA expression increases with age in brain tissues and fibroblasts. A. Scatter plots showing the correlation of TERRA v.s. age in brain, ovary and heart tissues. Each dot indicates the total TERRA level from all chromosome ends in an individual. P values, by Pearson’s correlation. B. TERRA levels in various tissues with different ages. Adult (21∼59 years); old (≥60 years). Bars, median with interquartile. P values, by Mann-Whitney U test. C. TERRA levels in human fibroblasts derived from healthy individuals of different ages. P values, by Pearson’s correlation. D. Boxplots showing the upregulation of TERRA levels in old human fibroblasts. Young (<30 years); adult (30∼59 years); old (≥60 years). Bars, median with interquartile. P values, by Mann-Whitney U test. E. Fibroblasts derived from HGPS patients display abnormal TERRA levels. Each dot represents the TERRA level of each individual. Bars, median with interquartile. P values, by Kolmogorov-Smirnov test.

### Elevated TERRA levels in aged fibroblasts and abnormal TERRA profiles in Hutchinson-Gilford progeria syndrome patients

Next, we quantified TERRA expression in fibroblasts obtained from healthy individuals of various ages and patients diagnosed with Hutchinson-Gilford progeria syndrome (HGPS), a genetic disorder characterized by premature aging features in childhood (56). Total TERRA levels, analyzed by TERRA-QUANT using poly(A)-enriched RNA-seq datasets, displayed a significant upregulation in fibroblasts derived from healthy older individuals (**Figures 6C-6D**), with an exponential increase observed in those over 75 years old. Chromosome-end-specific analysis showed that TERRA increased at multiple chromosome ends in the old group (**Supplementary Figure S8A**). Type I TERRA transcription was dominant in fibroblasts compared to Type II and III (**Supplementary Figure S8B**). Considering that HGPS patients were below 30 years old, we compared TERRA levels with those of their age-matched controls. Notably, HGPS patients displayed an aberrant and diverse TERRA expression pattern compared to healthy individuals (**Figure 6E**).

### Single-cell RNA-seq analysis reveals high TERRA levels in neurons

We assessed TERRA levels across various cell types by analyzing published RNA-seq datasets from human tissues, including adipose, blood, bone, brain, heart, lung, ovary, pancreas, and retina (**Supplementary Table 7**). Among these tissues, TERRA levels were particularly high in blood, bone, and ovary (**Figure 7A**). Given the highest level in blood cells, we sought to identify the specific cell types exhibiting high TERRA expression. To achieve this, we analyzed single-cell RNA-seq datasets (59) from peripheral blood mononuclear cells (PBMC) obtained from healthy young individuals. Notably, dendritic cells displayed the highest TERRA level among various immune cells (**Figure 7B**).

**Figure 7.**
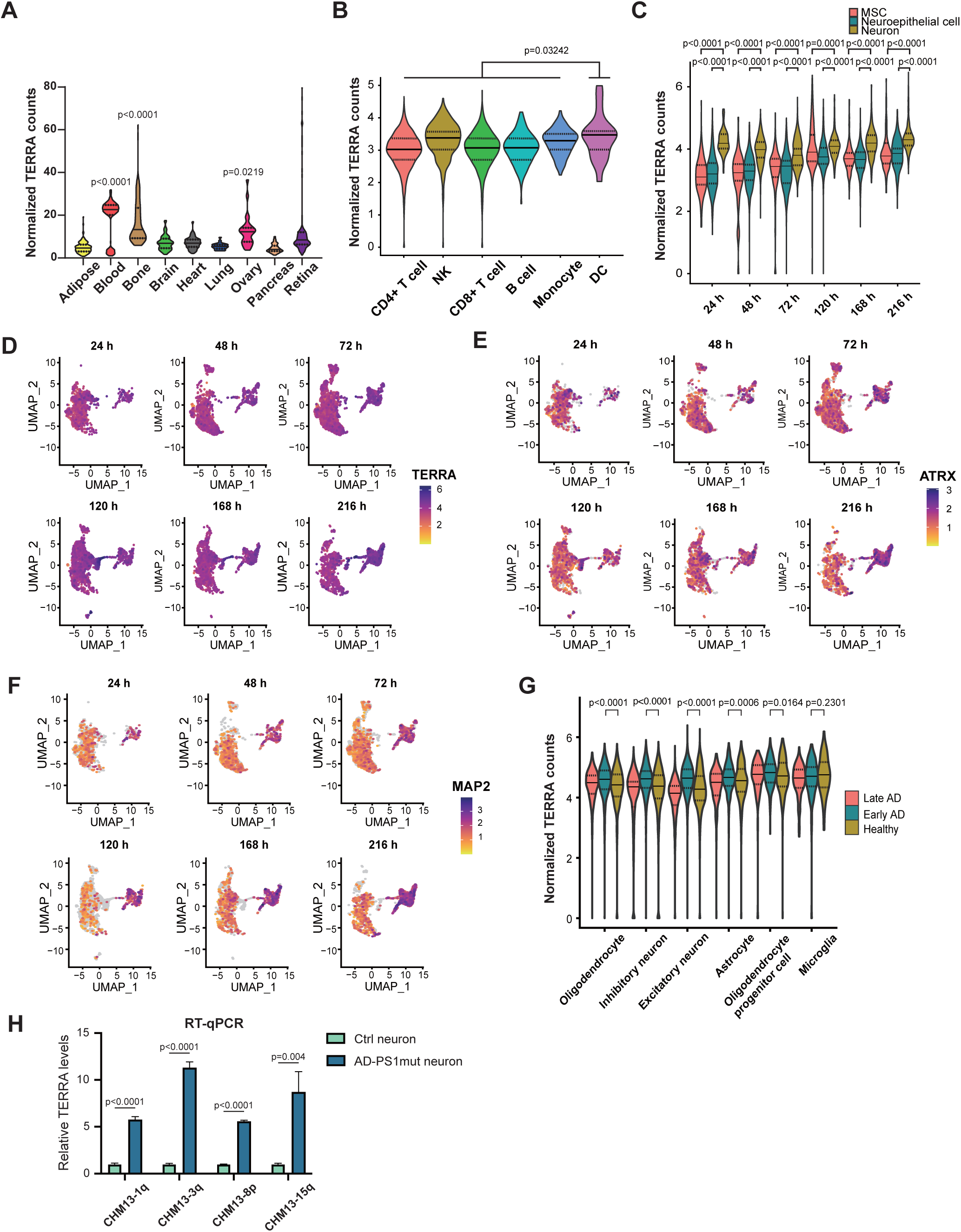
Single-cell analysis of TERRA levels undergoing neuronal differentiation and in Alzheimer’s disease. A. Violin plot showing normalized TERRA counts in various human tissues. The solid lines indicate the median and the dashed lines are interquartile range. P values, by Mann-Whitney U test. B. Single-cell RNA-seq showing normalized TERRA counts in various immune cells from PBMC. NK, nature killer cell; DC, dendritic cells. Solid lines, median. Dashed lines are interquartile range. P values, by Mann-Whitney U test. C. Single-cell RNA-seq analysis of normalized TERRA counts in human embryonic stem cells (hESCs) undergoing neuronal differentiation with various HOX patterning periods (24h,72h,48h, 120h, 168h, 216h). MSC, mesenchymal stem cells. Solid lines, median. Dashed lines are interquartile range. P values, by Mann-Whitney U test. D-F. Uniform manifold approximation and projection (UMAP) plots of single-cell RNA-seq datasets from hESCs undergoing neuronal differentiation using various HOX patterning periods. Colors according to TERRA, ATRX, or neuron marker MAP2 expression levels. G. TERRA levels in cells isolated from the prefrontal cortex of individuals with Alzheimer’s disease (AD). Single-nucleus RNA-seq datasets were obtained from the ROSMAP project, and grouped into healthy (no AD pathology), early-AD pathology, and late-AD pathology. Solid lines, median. Dashed lines are interquartile range. P values, by Mann-Whitney U test. H. RT-qPCR analysis showing elevated TERRA levels in AD neurons. AD-PS1mut or Ctrl neurons were differentiated from iPSC. AD-PS1mut carrying P117L mutation in *PS1* gene. Ctrl: mutation-corrected isogenic control. P values, by two-tailed Student’s t-test. n=3. TERRA levels normalized to GAPDH.

To investigate the potential role of TERRA in neurons, we examined TERRA levels using single-cell RNA-seq datasets obtained from human embryonic stem cells (hESCs) undergoing neuronal differentiation (57). The hESCs were converted to neuromesodermal progenitors (NMPs) with distinct HOX profiles along rostrocaudal axis by modulation of Wnt, FGF and retinoic acid signaling, and were further differentiated into ventral neuron phenotypes. Six different NMP cultures from hESCs were generated corresponding to HOX patterning periods: 24, 48, 72, 120, 168, and 216 hours. The cultures exhibited a progressive expression of caudal HOX paralogs, corresponding to cervical (HOX1-8; observed in 24h,48h, and 72h), thoracic (HOX1-9; observed in 120h), lumbar (HOX1-11; observed in 168h), and lumbosacral spinal regions (HOX1-13; observed in 216h). Cells were clustered based on gene expression into mesenchymal cells, neuroepithelial cells, and neurons. Notably, higher levels of TERRA were observed in neurons compared to neuroepithelial cells and mesenchymal cells (**Figure 7C**). Additionally, TERRA levels displayed a concordant expression pattern with ATRX during neuronal differentiation (**Figures 7D-7F**).

### Elevated TERRA levels during the early stage of Alzheimer’s disease progression

Next, we sought to analyze TERRA levels using TERRA-QUANT in neuronal disorder diseases. We analyzed published single-nucleus RNA-seq datasets from the prefrontal cortex of patients with Alzheimer’s disease (AD) (58). Postmortem human brain samples were obtained from participants enrolled in the Religious Order Study and Rush Memory and Aging Project (ROSMAP) (59). Individuals with AD pathology were categorized into two subgroups according to the previous study (58), reflecting distinct stages of AD pathological progression: 1) early-pathology characterized by amyloid burden, modest neurofibrillary tangles, and cognitive impairment; and 2) late-pathology marked by higher amyloid levels, elevated neurofibrillary tangles, increased global pathology, and cognitive impairment. Notably, increased TERRA levels were observed in various types of brain cells in the early stage of AD pathology compared to no AD pathology, with more profound TERRA levels in excitatory neurons (**Figure 7G, Supplementary Figure S9A**). To further investigate TERRA in AD neurons, induced pluripotent stem cells (iPSCs) derived from an AD patient, along with mutation-corrected isogenic control cells, were generated. The AD patient carried the P117L mutation in presenilin-1 (PS1mut), the catalytic subunit of γ-secretase. Notably, RT-qPCR demonstrated that AD-PS1mut neuronal cells differentiated from iPSCs exhibited significantly higher TERRA levels compared to control cells (**Figure 7H**), further supporting the upregulation of TERRA expression in AD neurons.

## Discussion

In this study, we investigated TERRA transcripts in human cells using Nanopore direct RNA-seq and TERRA capture RNA-seq. We identified TERRA transcription regions at 39 chromosome ends and utilized these defined regions to quantify TERRA expression across various tissues. Our bioinformatics pipeline is versatile, accommodating different RNA-seq datasets, including single-cell RNA-seq experiments. Notably, substantial mapped reads per cell were observed in single-cell RNA-seq datasets without YARN normalization (**Supplementary Figures S9A-S9C**), indicating that TERRA can be detected at the single-cell level.

Some discrepancies were observed between our findings and those reported by Rodrigues et al. (45) regarding TERRA expression at specific chromosome ends in HeLa and U2OS cells. A comparison of our and their results revealed consistent identification of TERRA expression at 31 chromosome ends, while 3 chromosome ends were consistently found to lack detectable TERRA expression (**Supplementary Figure S9D**). Our analysis revealed 8 additional chromosome ends expressing TERRA that were not reported by Rodrigues et al., while their study identified 4 ends absent from our dataset. We attribute these differences primarily to the distinct mapping strategies applied. Rodrigues et al. used a limited reference consisting of 25 kb subtelomeric regions, whereas we utilized the entire genome for read alignment. When we applied 25 kb subtelomeric regions as a reference genome for mapping our data, the results aligned closely with their findings. These observations suggest that the choice of mapping strategy impacts the sensitivity and specificity of TERRA detection at individual chromosome ends.

Commonly available datasets largely comprise RNA samples enriched with poly-A tails. Nevertheless, a significant portion of TERRA molecules lack poly-A tails, which may limit the detection of TERRA using poly-A enriched RNA-seq data. To investigate potential biases toward specific chromosome ends in datasets derived from poly(A)-enriched RNA-seq libraries, we performed poly(A)+ and poly(A)-TERRA capture sequencing in U2OS and HeLa cells. Our analysis revealed that poly(A)+ TERRA transcripts may preferentially be produced from certain chromosome ends and vice versa, consistent with the previous study (7). This preference is conserved in both U2OS and HeLa cells. Although there were detectable preference in TERRA detection from specific chromosome ends using poly(A)-enriched RNA-seq datasets, we still observed TERRA originating from multiple chromosome ends in aged fibroblasts, despite this bias (**Supplementary Figure S8A**). For analysis of total TERRA levels in human tissues, we accumulated TERRA reads from all chromosome ends to enhance read counts, thereby increasing overall TERRA counts for quantifying total TERRA levels. Due to low read counts mapped to individual chromosome ends, analyzing chromosome-end-specific TERRA levels remains challenging using publicly available RNA-seq datasets from human tissues. TERRA-enriched sequencing and longer read lengths could enable more precise quantification of TERRA at specific chromosome ends.

Utilizing TERRA-QUANT for quantifying TERRA levels across diverse tissues, we detected TERRA expression in most tissues examined. Notably, high TERRA levels were observed in blood and bone tissues, raising the possibility of a role for TERRA in immune-related processes. The earlier research has shown that TERRA can be secreted via extracellular exosomes to stimulate inflammatory cytokines during telomere dysfunction (60,61). TERRA also interacts with ZBP1 to activate immune response under conditions of replicative crisis (62). However, it is important to note that such ZBP1-TERRA complex-mediated immune activation is unlikely to occur under normal physiological conditions in healthy blood cells. Notably, we observed elevated total TERRA levels in fibroblasts, brain, and blood cells as individuals aged. The telomere shortening may contribute to TERRA upregulation during aging. However, neurons are postmitotic cells that are not expected to undergo telomere shortening. Therefore, the increase of TERRA in the brain is unlikely to result from telomere shortening, but may instead be associated with oxidative stress, neuronal damage, and telomere dysfunction in the aging brain. It has been previously reported that telomeric transcription is induced by DNA damage at telomeres in Hutchinson-Gilford progeria syndrome (HGPS) fibroblasts (33). In our analysis, fibroblasts derived from HGPS patients demonstrated abnormal TERRA expression, with some individuals showing high levels while others displayed lower levels. We reason that the diverse TERRA counts observed in HGPS patients could be attributed to variable telomere lengths or defects in telomere integrity. Previous studies have shown that telomere length in HGPS patients is generally shorter, yet exhibits variability (63,64). Low TERRA counts in some HGPS patients could result from either low TERRA expression or the loss of subtelomeric and telomeric DNA at chromosome ends.

Examination of single-nucleus RNA-seq datasets and RT-qPCR analysis of TERRA levels unveiled an upregulation of TERRA levels in Alzheimer’s disease (AD) neurons. AD is associated with several cellular stress responses—such as oxidative stress and chronic inflammation (65–67)—that are also known to upregulate TERRA expression (37,68). Moreover, the accumulation of DNA G-quadruplex (G4) structures has been reported in AD neurons (69), and TERRA has also been implicated in promoting G4 formation (27). These results indicate elevated TERRA levels correlate with increased DNA G4 formation and oxidative stress, which are aligned with the observations in AD neurons. Taken together, these connections provide a compelling rationale for further investigation into the role of TERRA in AD pathogenesis.

Upregulation of TERRA and ATRX during neuronal differentiation implies the potential function of TERRA in neuronal cells. Interestingly, the inverse relationship between TERRA and ATRX in the aged brain was observed, suggesting that low levels of ATRX might contribute to the dysfunction of the aged neurons. Previous research indicated that the absence of ATRX leads to cognitive impairment and promotion of cellular senescence (70), further highlighting the significance of exploring the interplay between TERRA, ATRX, and aging-related cellular processes in the brain and neuronal tissues. It is plausible that proper levels of TERRA and ATRX are crucial for maintaining normal gene expression in neuronal cells, and dysregulation of TERRA and ATRX may impact neuron function during aging.

Our study provides the annotation of TERRA transcripts in the T2T-CHM13 genome reference and introduces the bioinformatics tool “TERRA-QUANT” for quantifying TERRA levels. This lays the foundation for studying TERRA in diverse conditions, cell types, and human diseases.

## Data availability

TERRA capture RNA-seq datasets including Illumina short-read RNA-seq, Nanopore direct RNA-seq, poly(A)+ and poly(A)-TERRA capture RNA-seq, and the sequencing data of TERRA RT-qPCR products are accessible at GEO (GEO accession GSE250303) https://www.ncbi.nlm.nih.gov/geo/query/acc.cgi?acc=GSE250303. The published sequencing datasets analyzed in this study are described in **Supplementary Tables 7, 8, and 9**.

## Code availability

A copy of the custom code utilized for quantification of TERRA is accessible on GitHub (https://github.com/ls807terra/TERRA_RNA-seq_pipeline). Publicly available packages were used in this study as indicated in the Methods section.

## Funding

This work was supported by National Science and Technology Council (NSTC) grants NSTC 112-2628-B-002-008 (H-P. C. C.), NSTC 112-2320-B-002-058 (H-P. C. C.), NSTC 113-2320-B-002-009 (H-P. C. C.), NSTC 113-2628-B-002-010-MY3 (H-P. C. C.) and National Taiwan University grants NTU-111L7880 (H-P. C. C.), NTU-AS-112L104312 (H-P. C. C), NTU-CDP-112L7721 (H-P. C. C.), NTU-CDP-113L7705 (H-P. C. C.), and National Health Research Institutes grants NHRI-EX111-11107SI, NHRI-EX112-11107SI (H-P. C. C.), NHRI-EX113-11107SI (H-P. C. C.), NHRI-EX114-11411SI.

## Conflict of Interest declaration

The authors declare that they have no competing interests.

## Acknowledgments

We appreciated the insightful discussions with Liuh-Yow Chen, Ching-Shyi Peter Wu, as well as the contributions of our lab members Hong-Jhin Shen, Chia-Yu Guh, and Liv Weichien Chen. We thank Kevin, Tsai at IBMS, Academia Sinica, for his support with Nanopore Direct RNA-seq. We thank Technology Commons, College of Life Science, National Taiwan University for supporting the usage of equipment, and National Center for High-performance Computing (NCHC) of the National Institutes of Applied Research of Taiwan (NIAR) for providing computational and storage resources.

## Author contributions

H-P. C. C. and Y-H. H. conceived of and designed the study. Y-H. H. conducted TERRA capture, library constructions for sequencing, and data analysis. C-H. T., M-T. Y. P-C. Y, analyzed RNA-seq datasets. Y-C. C. performed qPCR for blood cells. C-P. Y. prepared samples for experiments. H-J. S. analyzed TERRA levels in AD neurons. C-H. Y and H-C. K contributed to AD iPSC generation and neuron differentiation. D-S H. collected blood samples. H-P. C. C., Y-H. H., C-H. T., P-C. Y, and M-T. Y. wrote the manuscript.

## Notes

### Competing Interest Statement

The authors have declared no competing interest.

### Summary of Updates

Including new data and the revised main text.

